# Dopamine ameliorates hyperglycemic memory-induced vascular dysfunction in diabetic retinopathy

**DOI:** 10.1101/2022.01.19.476943

**Authors:** Yeon-Ju Lee, Hye-Yoon Jeon, Ah-Jun Lee, Minsoo Kim, Kwon-Soo Ha

## Abstract

Dopamine is a neurotransmitter that mediates visual function in the retina, and hyperglycemic memory (HGM) is a pivotal phenomenon in the development of diabetic retinopathy (DR); however, the role of dopamine in HGM-induced retinal vascular dysfunction remains unclear. Here, we report a mechanism of HGM-induced retinal vascular dysfunction and the protective effect of dopamine against the HGM-induced DR. HGM induces retinal vascular lesions through persistent oxidative stress, mitochondrial dysfunction, and microvascular abnormalities after blood glucose normalization, and dopamine ameliorates this HGM-induced retinopathy. HGM induced persistent oxidative stress, mitochondrial membrane potential collapse and fission, and adherens junction disassembly and subsequent vascular leakage in the mouse retinas. These persistent hyperglycemic stresses were inhibited by dopamine treatment in human retinal endothelial cells and by intravitreal injection of levodopa in the retinas of HGM mice. Our findings suggest that dopamine alleviates HGM-induced retinal vascular dysfunction by inhibiting persistent mitochondrial dysfunction and microvascular abnormalities.

## INTRODUCTION

Dopamine, an important neurotransmitter in the nervous system, is synthesized from tyrosine by tyrosine hydroxylase in dopaminergic neurons and released in accordance with a circadian clock (Popova, 2014). In the retina, dopamine is released from dopaminergic amacrine cells and is involved in the light adaptive process of the visual system (Mohana Devi et al., 2020). Dopamine affects several neuronal cells in the retina, including photoreceptors and horizontal and bipolar cells, through five G-protein-coupled receptors named D1 to D5 (Popova, 2014). Dopamine levels are reduced in the retinas of diabetic rodent models, and supplementation of dopamine in these animals prevents visual defects (Aung et al., 2014; Chesler et al., 2021). In type 1 diabetic mice, dopamine deficiency contributes to early visual dysfunction, assessed by changes in spatial frequency threshold and contrast sensitivity, which is improved by intraperitoneal treatment with dopamine receptor agonists or the dopamine precursor levodopa (L-dopa) (Aung et al., 2014; Kim et al., 2018). A preclinical study in patients with diabetes revealed that early retinal dysfunction assessed by electroretinography is detectable prior to clinically recognized retinopathy and can be reversed by L-dopa treatment (Motz et al., 2020). Dopamine is therefore likely to be beneficial for hyperglycemia-induced early visual dysfunction in diabetes; however, its effect on retinal vascular dysfunction and protective mechanism remain unknown.

Diabetic retinopathy (DR) is the most common microvascular complication of diabetes and the leading cause of blindness in working-age populations (Wang and Lo, 2018). DR, a progressive metabolic disease from a non-proliferative stage to a proliferative stage, is caused by several risk factors including poor glycemic control, duration of diabetes, and hypertension (Kowluru, 2017; Simo-Servat et al., 2019); however, the treatment of DR remains challenging (Semeraro et al., 2019; Wang and Lo, 2018). Clinical trials showed that blood glucose normalization in patients with type 1 or type 2 diabetes does not arrest DR progression caused by persistent hyperglycemic stress, which is commonly referred to as hyperglycemic memory (HGM) or metabolic memory (Holman et al., 2008; Pirola et al., 2010; Zhang et al., 2012). The Diabetes Control and Complications Trial (DCCT) and the following up Epidemiology of Diabetes Interventions and Complications (EDIC) studies demonstrated that poor glycemic control leads to the development of diabetic complications, including DR, long after intensive glycemic control is achieved in patients with type 1 diabetes (Pirola et al., 2010). The United Kingdom Prospective Diabetes Study (USPDS) showed the long-term effects of intensive glycemic control on vitreous hemorrhage and retinal photocoagulation in patients with type 2 diabetes (Holman et al., 2008). The role of HGM in the development and progression of DR was also demonstrated in diabetic animal models, including diabetic dogs and rats (Engerman and Kern, 1987; Kowluru, 2017; Kowluru and Mohammad, 2020). It is now clear that HGM is the pivotal phenomenon in the development of DR, so exploration of the pathological mechanisms of HGM is needed to facilitate the development of new therapies for DR.

The underlying mechanisms of HGM have been investigated to uncover the pathophysiology of diabetic microvascular and macrovascular complications. Reactive oxygen species (ROS) generation is critically involved in the persistent hyperglycemic stress caused by HGM in endothelial cells and experimental animals (Ihnat et al., 2007; Lee et al., 2019; Paneni et al., 2013). Hyperglycemia induced upregulation of the pro-oxidant enzymes protein kinase C (PKC) β and NADPH oxidase subunit p47phox that persisted after restoration of normoglycemia in the retinas of diabetic rats (Ihnat et al., 2007). In the aortas of insulin-supplemented diabetic mice, activation of the mitochondrial adaptor protein p66^Shc^ by PKC βΙΙ persisted after return to normoglycemia, and the persistent p66^Shc^ activation was associated with continued ROS production and PKC βII upregulation, leading to a vicious cycle (Di Lisa et al., 2017; Paneni et al., 2012; Paneni et al., 2013). Sustained ROS generation upregulates expression of NF-kB subunit p65 and monomethylation of histone H3 at lysine 4 by the methyltransferase Set7/9, leading to overexpression of inflammatory genes including vascular cell adhesion molecule-1 and monocyte chemoattractant protein 1 (Okabe et al., 2012; Paneni et al., 2013). In the aortas of insulin-supplemented diabetic mice, persistent ROS generation formed a vicious cycle with transglutaminase 2 (TGase2), which plays an important role in HGM-induced expression of inflammatory adhesion molecules and apoptosis (Lee et al., 2019). Hence, an understanding of the persistent ROS generating machinery and its role in HGM-induced retinopathy is the real challenge in the development of effective therapies.

We hypothesized that dopamine ameliorates HGM-induced retinal vascular dysfunction in DR by inhibiting persistent mitochondrial dysfunction and microvascular abnormalities. We tested this in an HGM mouse model and found that intravitreal injection of L-dopa alleviated HGM-induced persistent oxidative stress, mitochondrial membrane potential (ΔΨ_m_) collapse and fission, and adherens junction disassembly and subsequent vascular leakage in the retina. Furthermore, we found that L-dopa supplementation mitigated HGM-induced pericyte degeneration, acellular capillary and pericyte ghost generation, and endothelial apoptosis. Our findings provide a useful window into the mechanism of HGM-induced retinal vascular dysfunction and suggest that dopamine might be a therapeutic agent for the treatment of DR.

## RESULTS

### Persistent hyperglycemic stress in the retinas of HGM mice

To investigate whether hyperglycemic stress persists in the retinas of HGM mice after blood glucose normalization, diabetic mice were exposed to hyperglycemic conditions for 12 weeks and then supplemented for 12 weeks with human recombinant insulin using osmotic pumps (**Fig. 1A**). Compared with non-diabetic controls, diabetic mice showed elevated water consumption, decreased body weight, and hyperglycemia, all of which were normalized by insulin supplementation (**Fig. 1B and C**). Hyperglycemia stimulated ROS generation in the retinas of diabetic mice; however, glucose normalization by insulin supplementation had no effect on the ROS generation in HGM mice (**Fig. 1D and E**). *In vivo* TGase activity was also elevated by hyperglycemia in the retinas of the diabetic mice, but the TGase activation was not affected by glucose normalization (**Fig. 1F**). Because ROS-mediated TGase activation is involved in vascular leakage in diabetic retinas (Lee et al., 2016), we studied vascular leakage in whole-mounted retinas. Hyperglycemia induced vascular leakage in the retinas of diabetic mice, and blood glucose normalization had no effect on the vascular leakage (**Fig. 1G and H**). These results suggest that oxidative stress, TGase activation, and vascular leakage are involved in persistent hyperglycemic stress in the retinas of HGM mice.

**Fig. 1.**
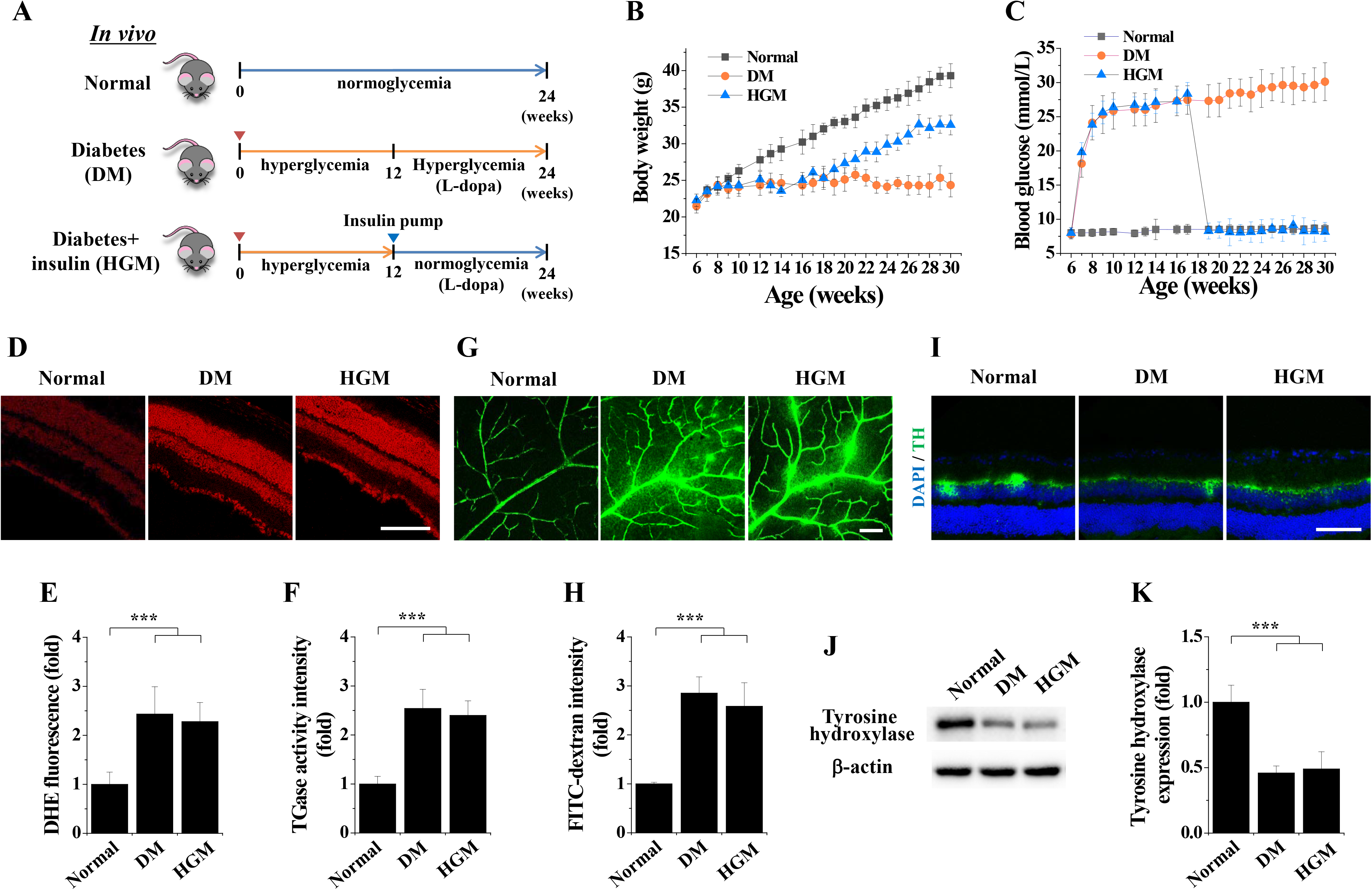
Persistent hyperglycemic stress and suppression of tyrosine hydroxylase expression in the retinas of diabetic mice supplemented with insulin. **A**: Schematic illustration of the generation of diabetic (DM) and insulin-supplemented diabetic (hyperglycemic memory; HGM) mice and administration of L-dopa. **B-K**: Twelve weeks after streptozotocin injection, diabetic mice were supplemented with insulin using osmotic pumps for 12 weeks. Reactive oxygen species (ROS) generation, vascular leakage, *in vivo* transglutaminase activity, and expression of tyrosine hydroxylase were visualized by confocal microscopy and quantitated in retinas from normal, diabetic, and HGM mice. B and C: Body weight (**B**) and blood glucose levels (**C**) were measured weekly (n = 10). D and **E**: ROS generation was visualized using dihydroethidium in retinal sections (**D**) and quantitated (**E**) by fluorescence intensity (n = 6). Scale bar, 100 μm. **F**: *In vivo* TGase activity in whole-mounted retinas (n = 6). **G** and **H**: Vascular leakage was visualized using FITC-conjugated dextran by fluorescence angiography in whole-mounted retinas (**G**) and quantitated (**H**) by fluorescence intensity (n = 6). Scale bar, 100 μm. **I-K**: Persistent suppression of tyrosine hydroxylase in retinal sections of HGM mice. **I**: Tyrosine hydroxylase expression was visualized by immunostaining (green) and nuclear counterstaining using Hochest 33342 (blue) in the retinal sections. Scale bar, 100 μm. **J** and **K**: Tyrosine expression was analyzed by Western blot (**J**) and quantitated by densitometry (n = 3) (**K**). \*\*\**P* < 0.001.

We next investigated whether tyrosine hydroxylase, the dopaminergic neuronal marker in retinal amacrine cells, is associated with persistent hyperglycemic stress in the retinal sections of HGM mice. Hyperglycemia reduced the expression of tyrosine hydroxylase in the retinas of diabetic mice, and the reduced expression was not recovered by glucose normalization in HGM mice (**Fig. 1I**). Persistent reduction of tyrosine hydroxylase expression in the retinas of HGM mice was further demonstrated by Western blot (**Fig. 1J and K**), suggesting that reduced dopamine levels are associated with the persistent hyperglycemic stress induced by HGM in mouse retinas.

### Inhibitory effects of dopamine on HGM-induced persistent oxidative stress and mitochondrial dysfunction in HRECs and mouse retinas

To study the effect of dopamine on persistent hyperglycemic stress, we first tested whether dopamine can inhibit sustained oxidative stress by measuring intracellular and mitochondrial ROS levels in HRECs treated with HGM conditions (**Fig. 2A**). High glucose conditions led to elevated intracellular ROS levels, which were sustained after glucose normalization but were inhibited by dopamine treatment (**Fig. 2B and C**). Similarly, elevated mitochondrial ROS levels were maintained in HRECs under HGM conditions but were suppressed by dopamine treatment (**Fig. 2D and E**). Dopamine also suppressed intracellular and mitochondrial ROS generation in HRECs treated with high glucose conditions. To validate our *in vitro* studies, we studied the role of dopamine in retinal sections of HGM mice supplemented with L-dopa for 12 weeks. Hyperglycemia led to elevated mitochondrial ROS levels in the retinas of diabetic mice, and the elevated mitochondrial ROS levels were sustained after blood glucose normalization but attenuated by L-dopa supplementation (**Fig. 2F and G**). These results indicate that dopamine inhibits HGM-induced persistent intracellular and mitochondrial ROS generation in HRECs and mouse retinas.

**Fig 2.**
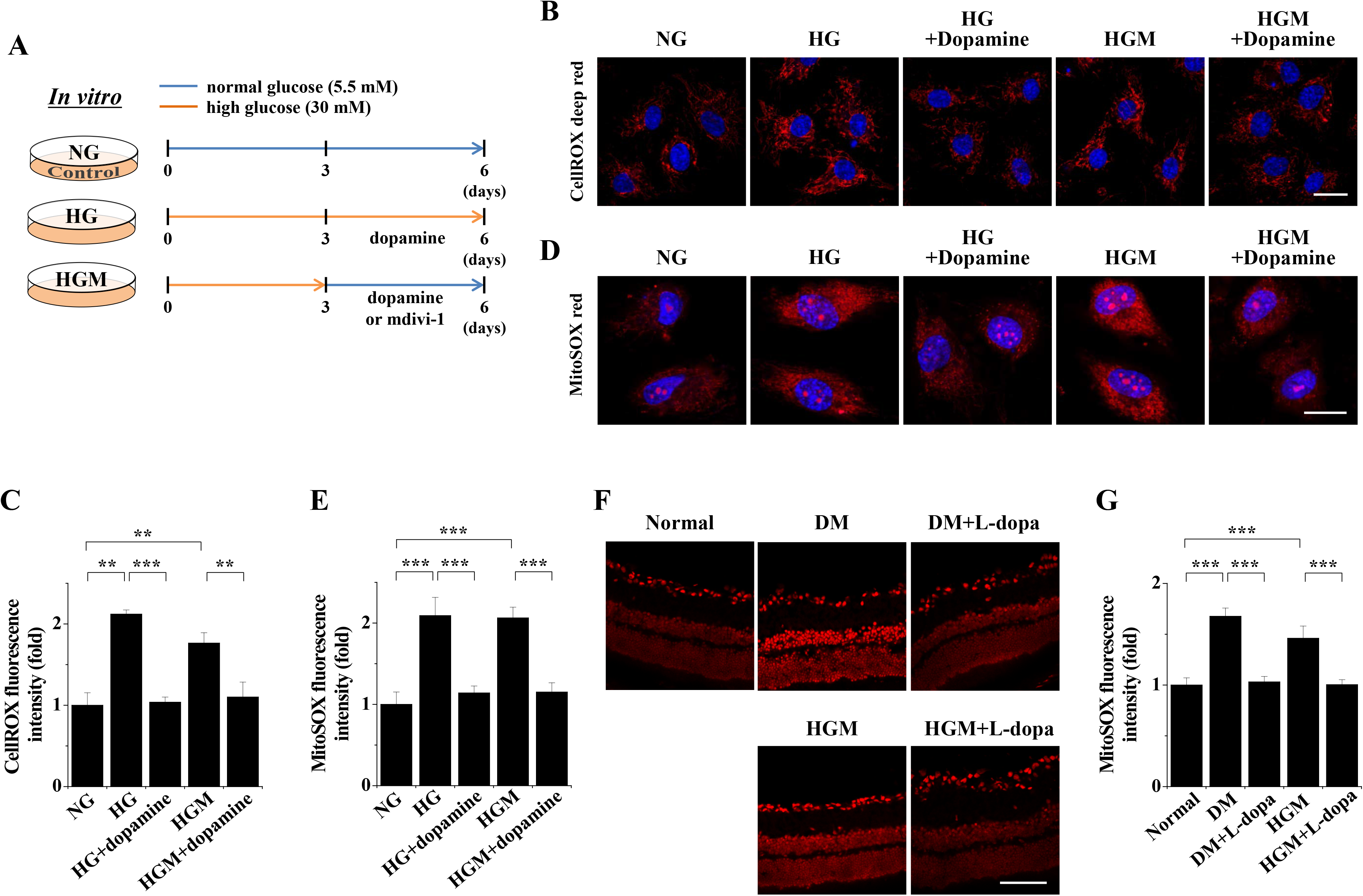
Dopamine inhibits persistent oxidative stress induced by HGM in human retinal endothelial cells (HRECs) and the retinas of HGM mice. **A:** Schematic illustration of the treatment of HRECs with dopamine in the high glucose and HGM conditions or with mdivi-1 in the HGM condition. **B-E:** HRECs were treated for 6 days with normal glucose (NG), high glucose (HG), high glucose in the presence of dopamine for the last 3 days (HG+dopamine), or high glucose for 3 days followed by normal glucose for 3 days (HGM) in the presence of 10 μM dopamine (HGM+dopamine). ROS generation was visualized and quantitated by confocal microscopy. **B** and **C:** Intracellular ROS generation was visualized by CellROX^TM^ deep red staining (red) and nuclear counterstaining using Hochest 33342 (blue) (**B**) and quantitated by fluorescence intensity (n = 3) (**C**). Scale bar, 25 μm. **D** and **E:** Mitochondrial ROS generation was visualized by MitoSOX^TM^ red staining (red) and nuclear counterstaining using Hochest 33342 (blue) (**D**) and quantitated by fluorescence intensity (n = 3) (**E**). Scale bar, 25 μm. **F** and **G:** Diabetic (DM) or insulin-supplemented diabetic (HGM) mice were supplemented with L-dopa (DM+L-dopa or HGM+L-dopa) for 12 weeks as illustrated in Fig. 1A. Mitochondrial ROS generation was visualized using MitoSOX^TM^ red in retinal sections (**F**) and quantitated by fluorescence intensity (n = 6) (**G**). Scale bar, 100 μm. \*\**P* < 0.01, \*\*\**P* < 0.001.

Because mitochondrial dysfunction is important in the development of DR (Kowluru, 2019), we investigated the inhibitory effect of dopamine on mitochondrial dysfunction by visualizing mitochondrial fission and ΔΨ_m_ collapse in HRECs. Exposure of HRECs to high glucose conditions induced mitochondrial fission and subsequent increase in the number of mitochondria per cell, which persisted after glucose normalization. These effects were inhibited by dopamine or mdivi-1, a potent inhibitor of Drp1-mediated mitochondrial fission (**Fig. 3A-C**). The ΔΨ_m_ collapse induced by the high-glucose treatment was sustained after glucose normalization and this sustained ΔΨ_m_ collapse was inhibited by dopamine or mdivi-1 treatment (**Fig. 3D and E**). We further explored the inhibitory effect of dopamine on HGM-induced mitochondrial dysfunction by Western blot (**Fig. 3F-H**). High-glucose-induced phosphorylation of Drp1 was sustained after glucose normalization, but the sustained phosphorylation was inhibited by dopamine. Dopamine also inhibited HGM-induced sustained expression of MFF, a key molecule that recruits Drp1 as a part of the mitochondrial fission machinery. Mdivi-1 had the same qualitative effects as dopamine on HGM-induced phosphorylation of Drp1 and expression of MFF. These results suggest that dopamine inhibits HGM-induced mitochondrial dysfunction by inhibiting sustained mitochondrial fission and ΔΨ_m_ collapse in HRECs.

**Fig 3.**
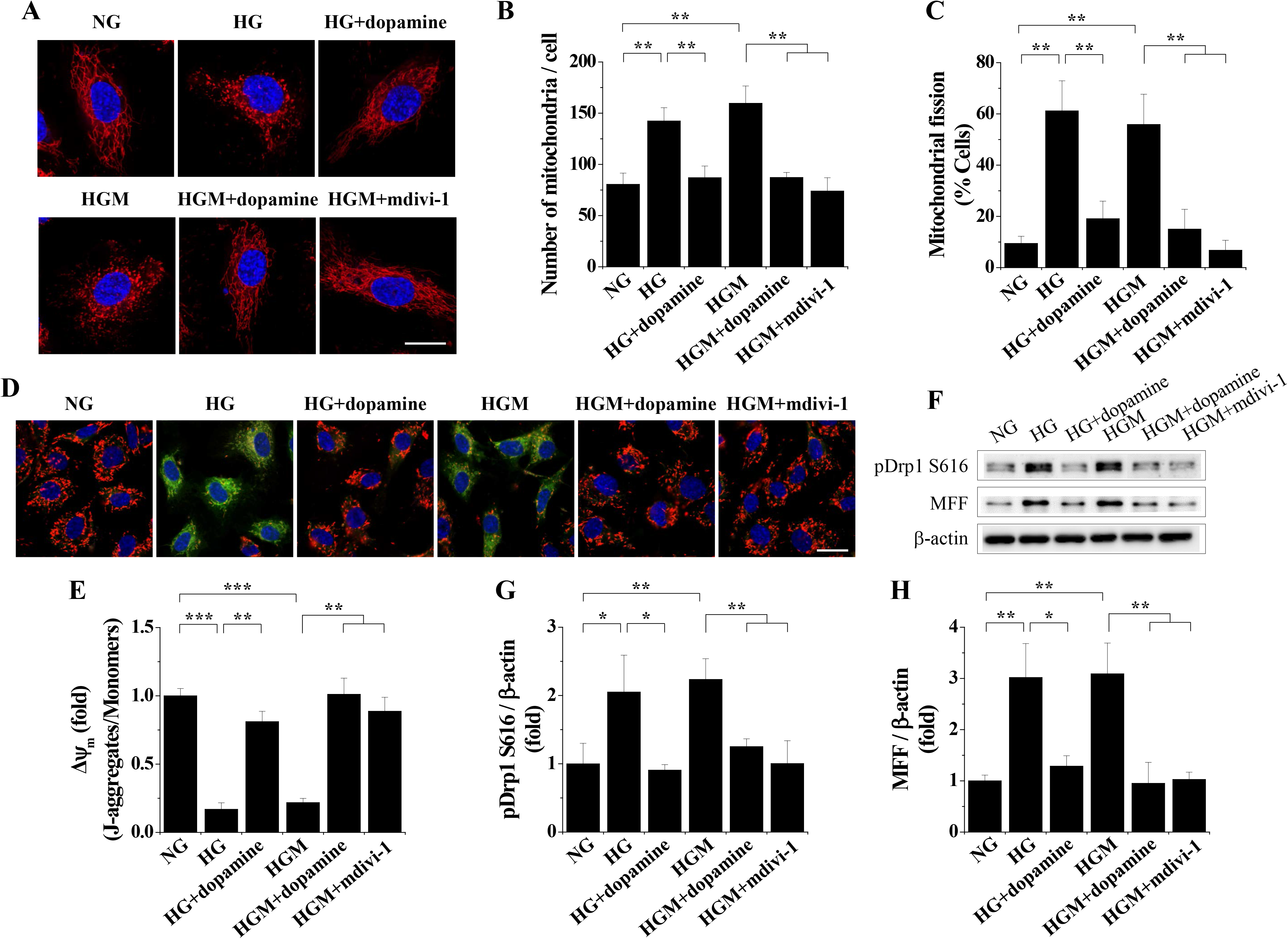
Dopamine inhibits persistent mitochondrial dysfunction induced by HGM in HRECs. HRECs were treated for 6 days with normal glucose (NG), high glucose (HG), high glucose in the presence of dopamine for the last 3 days (HG+dopamine), or high glucose for 3 days followed by normal glucose for 3 days (HGM) in the presence of 10 μM dopamine (HGM+dopamine) or 10 μM mdivi-1 (HGM+mdivi-1), as illustrated in Fig. 2A. Mitochondria, mitochondrial membrane potential (ΔѰ_m_), and TUNEL positive cells were visualized by confocal microscopy. Expression of mitochondrial dysfunction-related proteins was analyzed by Western blot. **A**-**C:** Mitochondria were visualized by MitoTracker Red CMXRos staining (red) and nuclear counterstaining using Hochest 33342 (blue) (**A**) and the number of mitochondria in cells was analyzed using ImageJ software (n = 3) (**B**). Scale bar, 25 μm. **C:** Mitochondrial fission was quantitated by confocal microscopy (n = 3). **D** and **E:** ΔΨ_m_ was visualized using JC-1 (**D**) and quantitated by measuring the J-aggregate (red) to monomer (green) fluorescence intensity ratio (n = 3) (**E**). Scale bar, 25 μm. **F-H:** Phosphorylation of dynamin-related protein 1 (Drp1) at Ser616 (pDrp1) and expression of mitochondrial fission factor (MFF) were analyzed by Western blot. **F:** Representative Western blot images. **G** and **H:** Densitometric quantitation of Drp1 phosphorylation (**G**) and MFF expression (**H**) using ImageJ software (normalized to β-actin levels) (n = 3). \**P* < 0.05, \*\**P* < 0.01, \*\*\**P* < 0.001.

To support our *in vitro* findings, we investigated the preventive effect of L-dopa supplementation on ΔΨ_m_ collapse, Drp1 phosphorylation, and MFF expression in the retinas of HGM mice. Hyperglycemia induced ΔΨ_m_ collapse in retinal sections from diabetic mice, and the ΔΨ_m_ collapse persisted after normoglycemia was improved by L-dopa treatment (**Fig. 4A and B**). Hyperglycemia-induced increases in Drp1 phosphorylation and MFF expression also persisted in the retinas of HGM mice after normoglycemia, but they were suppressed by L-dopa (**Fig. 4C-E**). Taken together, our results demonstrate that dopamine inhibits HGM-induced oxidative stress and mitochondrial dysfunction in HRECs and mouse retinas.

**Fig 4.**
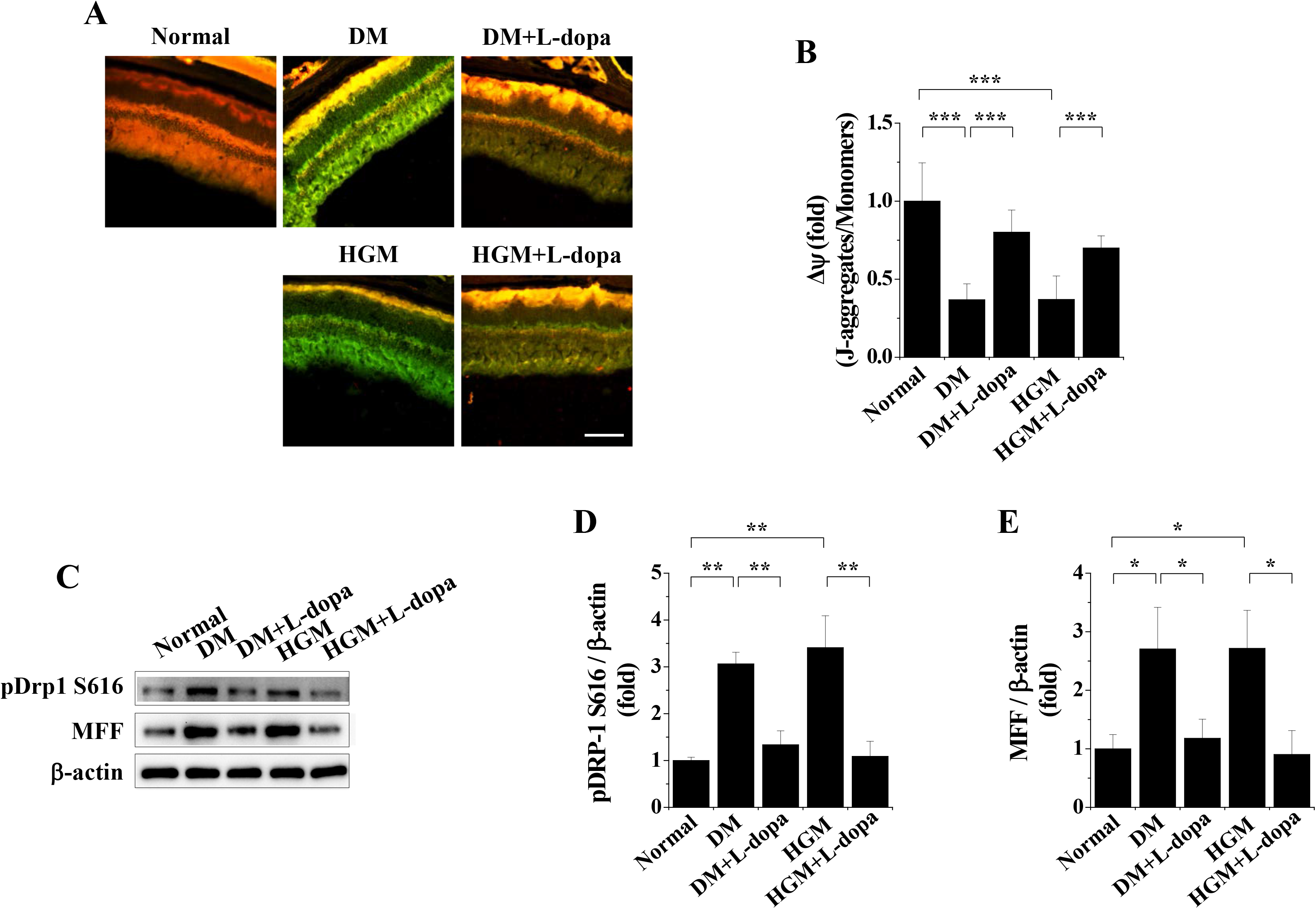
L-dopa inhibits HGM-induced ΔѰ_m_ collapse and expression of mitochondrial dysfunction-related proteins in mouse retinas. Twelve weeks after streptozotocin injection, diabetic (DM) mice were supplemented for 12 weeks with L-dopa (DM+L-dopa), insulin (HGM), or insulin and L-dopa (HGM+L-dopa), as illustrated in Fig. 1A. **A** and **B:** L-dopa inhibits persistent ΔѰ_m_ collapse in the retinas of HGM mice. ΔΨ_m_ was visualized by confocal microscopy using JC-1 in retinal sections from normal, DM, DM+L-dopa, HGM, and HGM+L-dopa mice (**A**) and quantitated by measuring the J-aggregate (red) to monomer (green) fluorescence intensity ratio (n = 8) (**B**). Scale bar, 100 μm. **C-E:** Phosphorylation of Drp1 at Ser616 (pDrp1) and expression of MFF were analyzed by Western blot in retinas from normal, DM, DM+L-dopa, HGM, and HGM+L-dopa mice. **C:** Representative Western blot images. **D** and **E:** Densitometric quantitation of Drp1 phosphorylation (**D**) and MFF expression (**E**) using ImageJ software (normalized to β-actin levels) (n = 3). \**P* < 0.05, \*\**P* < 0.01, \*\*\**P* < 0.001.

### Inhibitory effects of dopamine on HGM-induced adherens junction disassembly and vascular leakage in HRECs and mouse retinas

To understand the role of dopamine in vascular leakage, we investigated the inhibitory effects of dopamine on VEGF-induced adherens junction disassembly and vascular permeability in HRECs. Hyperglycemia elevated the levels of VEGF in the retinas of diabetic mice, and the elevated levels were sustained after blood glucose normalization in HGM mice (**Fig. 5A**), highlighting the important role of VEGF in diabetes-associated retinopathy. In HRECs, VEGF induced VE-cadherin disassembly and subsequent internalization of VE-cadherin, which were suppressed by dopamine (**Fig. 5B-E**). We investigated the role of dopamine in VEGF-induced vascular leakage by *in vitro* endothelial cell monolayer permeability assay in HRECs and *in vivo* Miles vascular permeability assay in mouse ears. In HRECs, VEGF caused an increase in endothelial cell permeability, which was reversed by dopamine (**Fig. 5F**). *In vivo* vascular leakage was elevated by intradermal injection of VEGF into mouse ears compared with that in PBS-injected ears, and the elevated vascular leakage was prevented by dopamine (**Fig. 5G and H**). These results indicate that dopamine prevents VEGF-induced adherens junction disassembly and subsequent increase in vascular permeability in HRECs and mouse ears.

**Fig 5.**
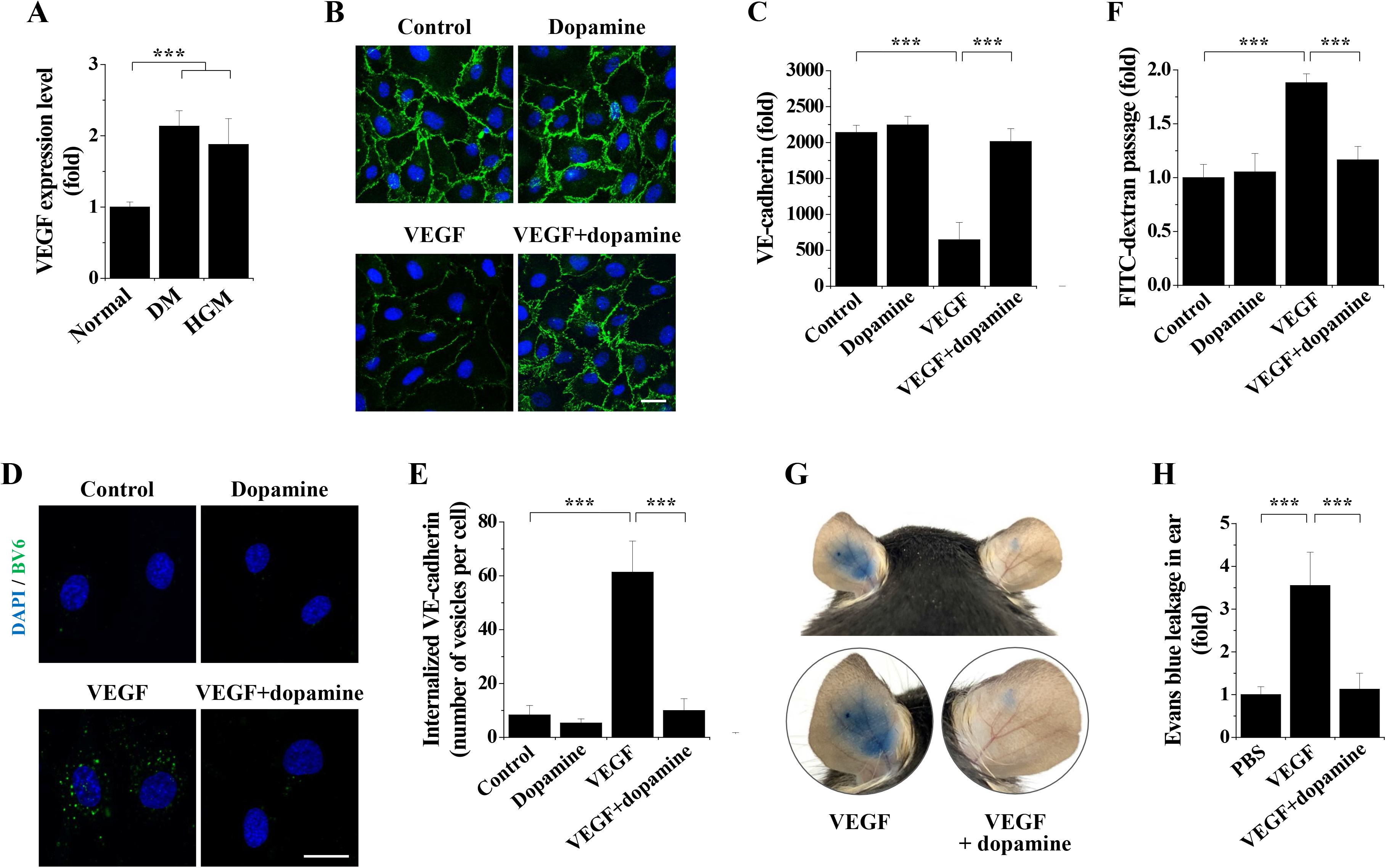
Dopamine inhibits VEGF-induced vascular leakage in HRECs and mouse ears. **A:** Analysis of VEGF expression levels in the retinas from normal, DM, and HGM mice (n = 6). **B-F:** HRECs were incubated with 10 μM dopamine for 30 min and treated with 10 ng/ml VEGF for 90 min. VE-cadherin (**B** and **C**) and internalization of VE-cadherin (**D** and **E**) were visualized by confocal microscopy. *In vitro* endothelial cell permeability was quantitated by spectrofluorometry (**F**). **B** and **C**: Visualization of VE-cadherin. VE-cadherin was visualized by immunofluorescence staining with nuclear counterstaining using DAPI (**B**) and adherens junctions were quantitated by measuring the fluorescence intensities (n = 3) (**C**). Scale bar = 25 μm. **D** and **E**: Internalization of VE-cadherin. Internalization of VE-cadherin was visualized by staining of cell surface VE-cadherin extracellular domain with nuclear counterstaining using DAPI (**D**) and quantitated by measuring the number of vesicles per cell (n = 3) (**E**). Scale bar = 25 μm. **F:** *In vitro* endothelial cell permeability was determined using 40-kDa FITC-dextran (n = 3). **G** and **H**: Miles vascular permeability assay in mouse ears. VEGF was injected intradermally into the middle part of mouse ears alone or in combination with dopamine (VEGF+dopamine). Extravasated Evans blue was photographed (**G**) and quantitated by spectrometry (n = 6) (**H**). \*\*\**P* < 0.001.

To support our findings in HRECs and mouse ears, we further studied the effect of dopamine on HGM-induced adherens junction disassembly and vascular leakage in mouse retinas. Compared with non-diabetic controls, the retinas of diabetic mice displayed hyperglycemia-induced VE-cadherin disassembly in the superficial and deep vascular layers, which persisted after return to normoglycemia (**Fig. 6A and B**). The HGM-induced disassembly of VE-cadherin was alleviated by L-dopa supplementation. High levels of FITC-dextran extravasation were sustained in the retinas of HGM mice after blood glucose normalization, but the HGM-induced vascular leakage was inhibited by L-dopa supplementation (**Fig. 6C and D**). We further studied the inhibitory effect of dopamine against HGM-induced retinal vascular leakage by Evans blue assay. Evans blue extravasation persisted after glucose normalization in the retinas of HGM mice but was suppressed by L-dopa (**Fig. 6E**). Together, our findings demonstrate that L-dopa ameliorates HGM-induced adherens junction disassembly and subsequent microvascular leakage in mouse retinas.

**Fig 6.**
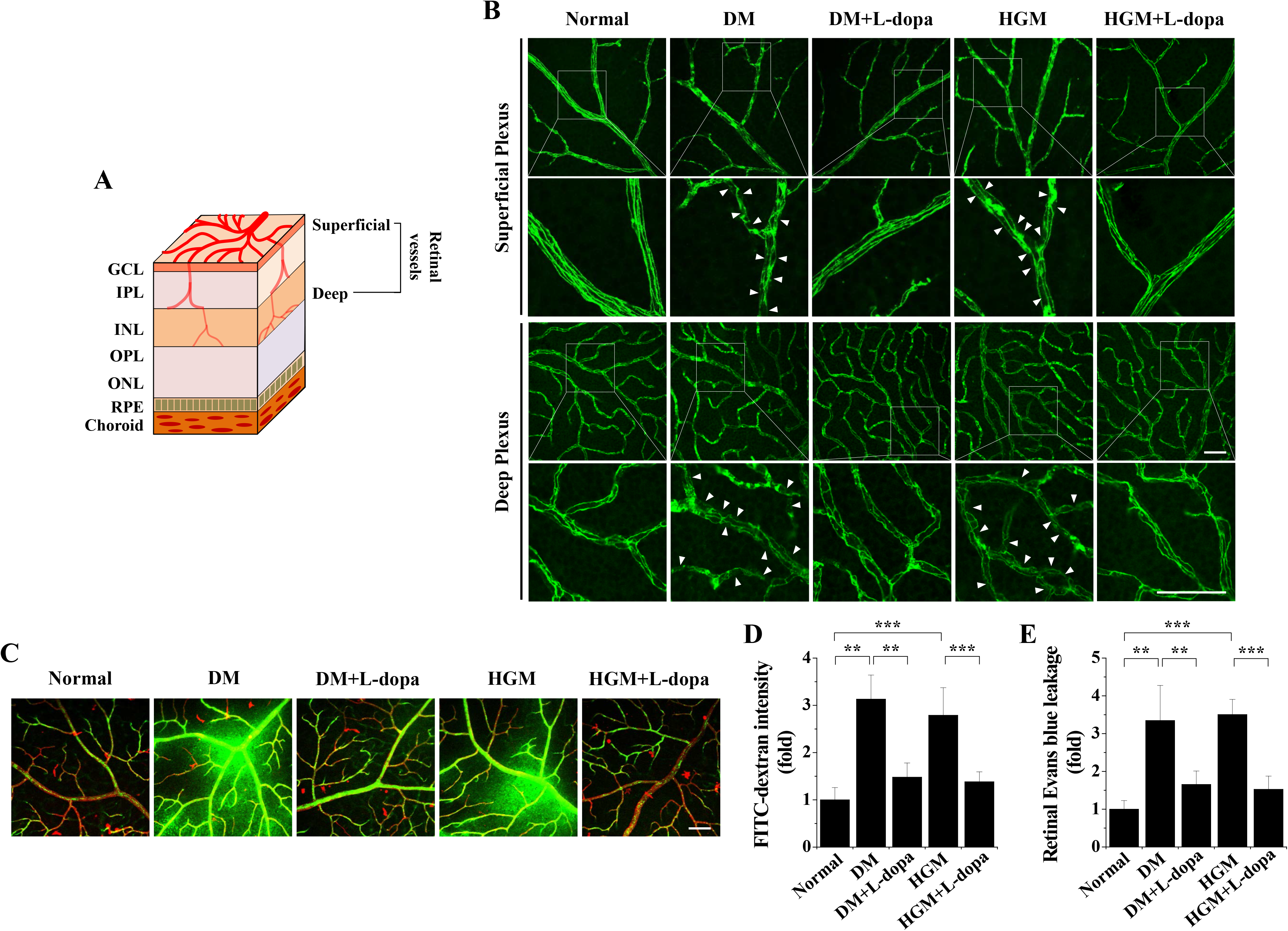
L-dopa prevents HGM-induced adherens junction disassembly and vascular leakage in mouse retinas. **A:** Schematic cross-sectional image of retinal neuronal and vascular structure. GCL, ganglion cell layer; IPL, inner plexiform layer; INL, inner nuclear layer; OPL, outer plexiform layer; ONL, outer nuclear layer; RPE, retinal pigment epithelium. **B:** VE-cadherin was visualized by confocal microscopy in the superficial and deep layers of whole-mounted retinal vessels from normal, diabetic (DM), L-dopa-supplemented (DM+L-dopa), HGM, and L-dopa-supplemented HGM (HGM+L-dopa) mice. The square areas in each image in the top row are displayed as magnified images in the bottom row. Arrows indicate disrupted adherens junctions. Scale bar, 50 μm. **C and D:** Vascular leakage was visualized using FITC-dextran (green) by fluorescence angiography in whole-mounted retinas from the five groups of mice (**C**) and quantified by the fluorescence intensity (n = 8) (**D**). Vessels were stained using Isolectin B4 (red). Scale bar, 100 μm. **E**: Quantitative analysis of Evans blue leakage in retinas (n = 8). \*\**P* < 0.01, \*\*\**P* < 0.001.

### Inhibitory effects of L-dopa on microvascular abnormalities and endothelial apoptosis in the retinas of HGM mice

To investigate whether microvascular abnormalities that represent retinopathy persist after normoglycemia, we analyzed pericyte degeneration and the numbers of acellular capillaries and pericyte ghosts in the retinas of HGM mice. Hyperglycemia led to a reduction in the numbers of NG2-positive pericytes in the retinas of diabetic mice, which persisted after blood glucose normalization in HGM mice but was reversed by L-dopa (**Fig. 7A and B**). The numbers of acellular capillaries and pericyte ghosts in trypsin-digested retinas of diabetic mice increased by hyperglycemia, and the increased numbers were maintained after blood glucose normalization but were reduced by L-dopa treatment (**Fig. 7C-E**). These results show that hyperglycemia-induced microvascular abnormalities in the mouse retina are sustained after blood glucose normalization but are ameliorated by L-dopa supplementation.

**Fig 7.**
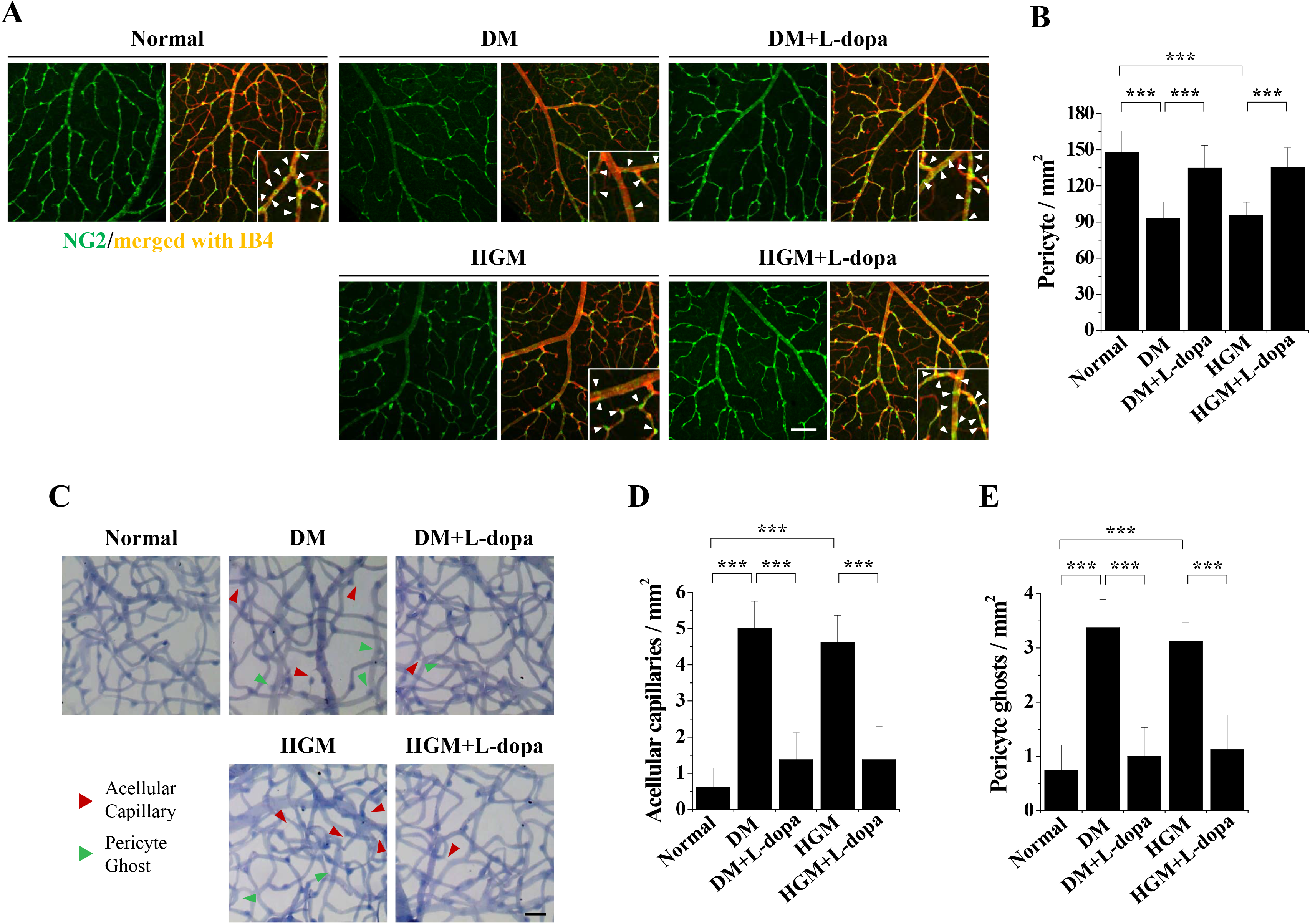
L-dopa prevents HGM-induced pericyte loss and altered microvascular integrity in mouse retinas. **A** and **B**: L-dopa prevents HGM-induced pericyte loss. **A:** Pericytes were visualized using NG-2 (green) by confocal microscopy with vessel staining using Isolectin B4 (IB4, red) in whole-mounted retinas from normal, diabetic (DM), L-dopa-supplemented (DM+L-dopa), HGM, and L-dopa-supplemented HGM (HGM+L-dopa) mice. Arrows in the magnified merged images indicate pericytes. Bar, 100 μm. **B**: Calculation of the pericyte numbers per mm^2^ of retinal vessels using ImageJ software (n = 8). **C-E:** L-dopa prevents HGM-induced formation of acellular capillaries and pericyte ghosts. Acellular capillaries (red arrows) and pericyte ghosts (green arrows) were visualized in whole-mounts of trypsin-digested retinas from the five groups of mice (**C**) and quantitated per mm^2^ vascular area (n = 8) (**D** and **E**). Scale bar, 100 μm. \*\*\**P* < 0.001.

We next studied the role of L-dopa in endothelial cell apoptosis by Western blot and TUNEL staining of the retinas of HGM mice. The expression of BAX and cytochrome c was maintained in the retinas of HGM mice after return to normoglycemia, but L-dopa treatment attenuated the HGM-induced expression of these proteins (**Fig. 8A-C**). Hyperglycemia increased the number of TUNEL-positive cells in diabetic retinas, and the HGM-induced increase in the number of apoptotic cells was normalized by L-dopa (**Fig. 8D and E**). These results suggest that L-dopa alleviates HGM-induced endothelial apoptosis in the mouse retina. Taken together, our results suggest that HGM causes vascular dysfunction through persistent mitochondrial dysfunction and microvascular abnormalities after blood glucose normalization in the mouse retina, and that dopamine ameliorates the HGM-induced retinal vascular dysfunction by reversing the mitochondrial dysfunction and vascular abnormalities (**Fig. 8F**).

**Fig 8.**
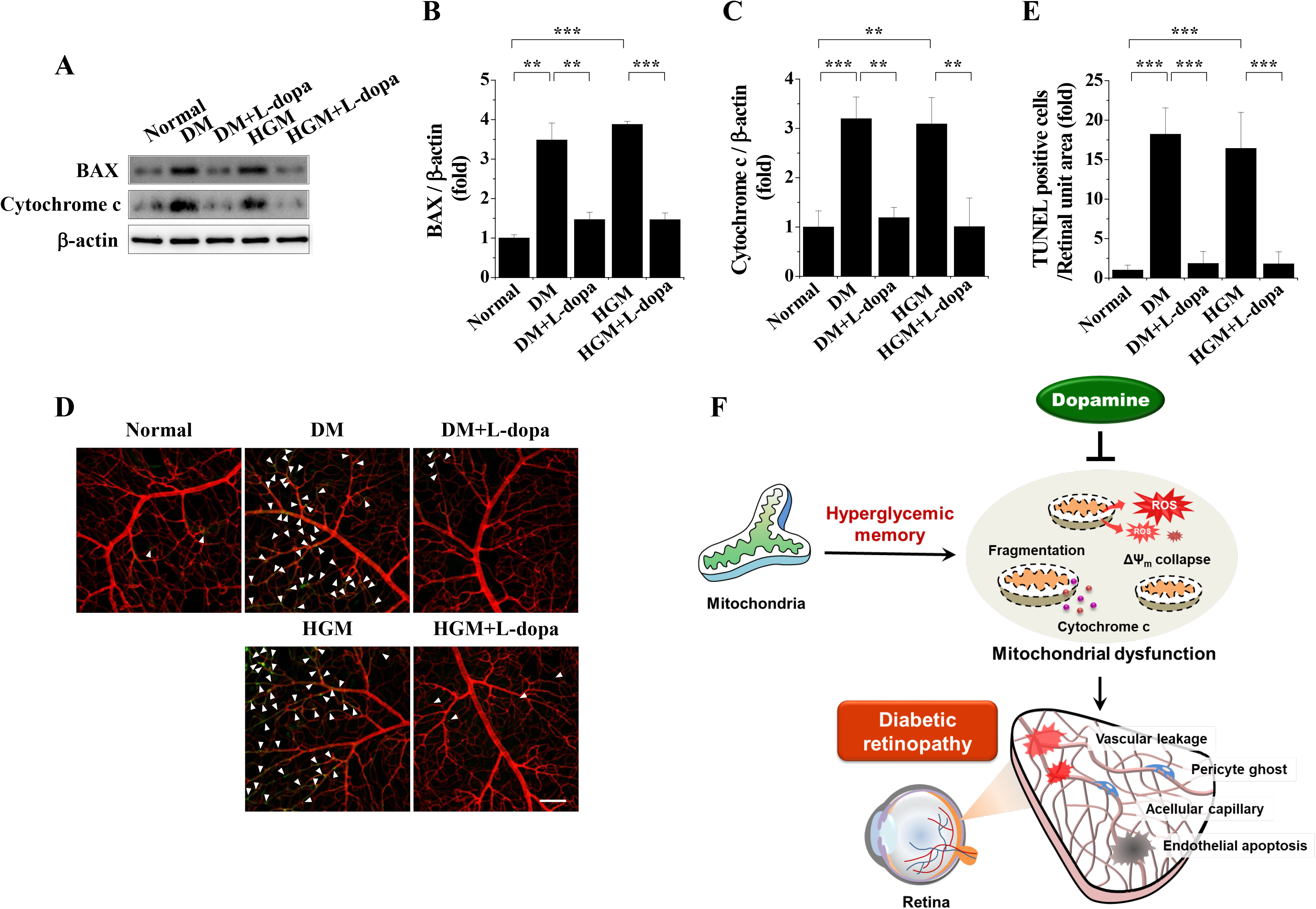
L-dopa prevents HGM-induced apoptosis of retinal vascular endothelial cells and a schematic diagram of the role of dopamine in HGM-induced mitochondrial dysfunction and diabetic retinopathy. **A-C:** Expression of BAX and cytochrome c were analyzed by Western blot in retinas from normal, DM, DM+L-dopa, HGM, and HGM+L-dopa mice. **A:** Representative Western blot images. **B** and **C**: Densitometric quantitation of BAX (**B**) and cytochrome c (**C**) expression using ImageJ software (normalized to β-actin levels) (n = 3). **D** and **E**: Apoptotic endothelial cells were stained by TUNEL with vessel counterstaining using Isolectin B4 in whole-mounts of trypsin-digested retinas (**D**) and quantitated by confocal microscopy (n = 8) (**E**). Scale bar, 100 μm. \*\**P* < 0.01, \*\*\**P* < 0.001. **F:** A model depicting the protective effects of dopamine against HGM-mediated mitochondrial dysfunction and microvascular abnormalities in diabetic mice.

## DISCUSSION

Dopamine is synthesized in the retina by tyrosine hydroxylase in dopaminergic amacrine neurons and is involved in visual signaling. Dopamine is beneficial for hyperglycemia-induced visual dysfunction in the retinas of diabetes; however, its role in HGM-induced vascular dysfunction remains unknown. HGM, or persistent hyperglycemic stress after glucose normalization, is a pivotal phenomenon in the development of DR. HGM-induced retinopathy was initially observed by Engerman and Kern (Engerman and Kern, 1987), who found that DR is not improved by good glycemic control in diabetic dogs. The DCCT-EDIC studies demonstrated that episodes of poor glycemic control contribute to the development of DR long after improved glucose control is achieved in patients with type 1 diabetes (Pirola et al., 2010). This long-lasting effect is also reported in patients with type 2 diabetes in the UKPDS study (Holman et al., 2008; Pirola et al., 2010). In the retinas of diabetic rats, oxidative stress and hypermethylation of mitochondrial proteins are not inhibited by reversal of hyperglycemia after poor glycemic control (Kowluru, 2003; Kowluru and Mohammad, 2020). Therefore, understanding the underlying mechanism of HGM is essential for the development of new therapies for HGM-induced retinopathy. In this study, we demonstrated that dopamine ameliorated HGM-induced retinal vascular dysfunction by normalizing persistent mitochondrial dysfunction and microvascular abnormalities. HGM induced persistent oxidative stress, mitochondrial fission and ΔΨ_m_ collapse, and subsequent adherens junction disassembly and microvascular leakage leading to retinopathy, as shown by increases in endothelial apoptosis and the numbers of acellular capillaries and pericyte ghosts. L-dopa ameliorated HGM-induced vascular dysfunction by inhibiting persistent mitochondrial dysfunction and microvascular abnormalities in the retina, suggesting dopamine as a possible therapeutic agent for HGM-induced retinopathy.

Mitochondria play a central role in the pathogenesis of DR (Kowluru, 2019). Oxidative stress induces mitochondrial dysfunction by impairing mitochondrial DNA stability and mitochondrial dynamics, causing subsequent mitochondrial fission (Kim et al., 2020; Kowluru, 2019). Mitochondrial fission through cytochrome c release promotes retinal vascular apoptosis (Kim et al., 2020; Wu et al., 2011); however, the role of mitochondrial dysfunction in HGM-induced retinopathy is unknown. We showed that in HRECs, HGM induced persistent mitochondrial ROS generation and subsequent mitochondrial dysfunction through sustained ΔΨ_m_ collapse, mitochondrial fission, and subsequent increase in the number of mitochondria after glucose normalization. In the retinas of diabetic mice, HGM consistently induced persistent mitochondrial ROS generation, ΔΨ_m_ collapse, and increases in Drp1 phosphorylation and MFF expression after return to normoglycemia. These results indicate that mitochondrial dysfunction contributes to the pathogenesis of HGM-induced vascular dysfunction and therefore might be an effective target for the treatment of DR.

Our results show that in addition to its role in hyperglycemia-induced neuronal dysfunction, dopamine ameliorates HGM-induced microvascular damage. Although DR is considered a diabetic microvascular complication, it is reported that neurodegeneration is also involved in DR pathogenesis, and thus DR is a neurovascular complication caused by microvascular dysfunction and neurodegeneration (Simo-Servat et al., 2019; Simo et al., 2018). Although dopamine deficiency contributes to early visual dysfunction in rodent models of diabetes, and this hyperglycemia-induced visual dysfunction is improved by dopamine treatment (Aung et al., 2014; Chesler et al., 2021; Kim et al., 2018), the role of dopamine in diabetes-associated retinal vascular damage has been unclear. Our results demonstrate that dopamine has a protective effect against persistent microvascular leakage and abnormalities induced by HGM. In HRECs, dopamine prevented VEGF-induced adherens junction disassembly and subsequent endothelial cell permeability. In the retinas of HGM mice, L-dopa supplementation suppressed HGM-induced adherens junction disassembly and subsequent microvascular leakage, and it ameliorated microvascular abnormalities including pericyte degeneration, acellular capillary and pericyte ghost generation, and endothelial apoptosis. Furthermore, L-dopa suppressed HGM-induced persistent mitochondrial ROS generation and ΔΨ_m_ collapse in retinal neuronal cells. Together, these results suggest that dopamine might ameliorate HGM-induced retinopathy by normalizing microvascular abnormality and neuronal dysfunction.

It is likely that dopamine ameliorates HGM-induced microvascular dysfunction in the retina by inhibiting VEGF-induced oxidative stress leading to vascular leakage. VEGF, a vascular permeability factor, plays a key role in DR (Lee et al., 2016). Previously, it was reported that VEGF activates transglutaminase 2 (TGase2) by intracellular Ca^2+^ elevation and ROS generation and subsequently induces microvascular leakage through adherens junction disassembly in the retinas of diabetic mice (Lee et al., 2017; Lee et al., 2018). We found that VEGF levels were persistently elevated in the retinas of HGM mice and that dopamine prevented vascular leakage induced by intradermal injection of VEGF into mouse ears. Consistent with our *in vivo* results, dopamine treatment of HRECs inhibited VEGF-induced intracellular events including intracellular and mitochondrial ROS generation, adherens junction disassembly, and subsequent endothelial cell permeability. It is reported that dopamine inhibits vascular permeability induced by VEGF in mice (Basu et al., 2001). In human umbilical vein endothelial cells, dopamine dephosphorylates VEGF receptor 2 by activating the Src-homology-2-domain-containing protein tyrosine phosphatase 2 through D2 dopamine receptors (Sinha et al., 2009). These findings suggest that VEGF-induced molecular events play a key role in the pathogenesis of HGM-induced retinopathy, and that dopamine ameliorates these events by regulating VEGF 2 receptors. It is still necessary, however, to elucidate the molecular mechanism of dopamine action in the retina.

In conclusion, HGM induces DR through persistent oxidative stress, mitochondrial dysfunction, and microvascular abnormalities in the retina after blood-glucose normalization. Dopamine alleviates HGM-induced vascular dysfunction by inhibiting the persistent mitochondrial dysfunction and microvascular abnormalities. Our results provide a possible mechanism of HGM-induced vascular dysfunction and suggest dopamine as a possible therapeutic agent for the treatment of DR.

## METHODS

### Cell culture

Human retinal endothelial cells (HRECs) were purchased from the Applied Cell Biology Research Institute (Cell Systems, Kirkland, WA, USA) and grown, as previously described (Lee et al., 2018). For experiments, endothelial cells from passages 4 to 7 were incubated for 12 h in low-serum medium supplemented with 2% fetal bovine serum, 0.1 ng/mL basic fibroblast growth factor, and antibiotics, and then subjected to treatments with 5.5 mM D-glucose for 6 days (normal glucose), 30 mM glucose for 6 days (high glucose), or 30 mM glucose for 3 days followed by 5.5 mM glucose for 3 days (HGM).

### Measurement of ROS levels in endothelial cells

Intracellular and mitochondrial ROS levels were measured using CellROX^TM^ deep red and MitoSOX^TM^ red mitochondrial superoxide indicator (Thermo Fisher Scientific, Waltham, MA, USA), respectively, as previously described (Bhatt et al., 2013). Fluorescence intensities of single stained cells were determined by confocal microscopy (K1-Fluo; Nanoscope Systems, Daejeon, Korea). The ROS levels were determined by comparing the average fluorescence intensities of treated cells with those of control cells (fold change).

### Analysis of mitochondrial fission and ΔΨ_m_ in endothelial cells

Mitochondrial fission was analyzed using MitoTracker Red CMXRos, as previously described (Bhatt et al., 2013). Briefly, endothelial cells were incubated at 37°C with 100 nmol/L MitoTracker Red CMXRos (Thermo Fisher Scientific) and 0.5 μg/mL Hochest 33342 (MilliporeSigma) for 30 min. Cells with disrupted and predominantly spherical mitochondria were identified as having mitochondrial fission. Thirty cells per experiment were used to calculate the percentage of cells undergoing mitochondrial fission.

ΔΨ_m_ was determined using 2 μmol/L JC-1 (Thermo Fisher Scientific) and confocal microscopy, as previously described (Bhatt et al., 2013). Data are expressed as the J-aggregate to monomer fluorescence intensity ratio.

### Visualization and internalization of VE-cadherin in endothelial cells

Endothelial cells were incubated with 10 μM dopamine for 30 min and stimulated with 10 ng/ml vascular endothelial growth factor (VEGF) for 90 min. Vascular endothelial (VE)-cadherin was visualized by confocal microscopy as previously described (Lee et al., 2016). Briefly, following fixation and permeabilization, endothelial cells were incubated overnight with a monoclonal VE-cadherin antibody (1:200; Santa Cruz Biotechnology, Dallas, TX, USA) at 4°C. The cells were then probed with a fluorescein isothiocyanate (FITC)-conjugated goat anti-mouse antibody (1:200; MilliporeSigma) for 2 h and stained with 1 μg/mL 4′,6-diamidino-2-phenylindole (DAPI; MilliporeSigma) for 10 min. VE-cadherin levels were represented by histograms and quantitatively analyzed using the peak fluorescence intensities of the histograms at the single-cell level.

Internalization of VE-cadherin was visualized by immunofluorescence as previously described (Lee et al., 2017). Briefly, the cells were incubated for 30 min at 4°C with a monoclonal antibody against the cell-surface VE-cadherin extracellular domain (BV6; MilliporeSigma) and then acid washed for 10 min with a low-pH buffer containing 100 mM glycine, 20 mM magnesium acetate, and 50 mM potassium chloride (pH 2.2). Following fixation and permeabilization, the cells were probed with an FITC-conjugated goat anti-mouse antibody for 2 h and stained with 1 μg/mL DAPI for 10 min.

### *In vitro* endothelial cell monolayer permeability assay

*In vitro* endothelial cell monolayer permeability assays were performed as previously described (Jeon et al., 2019). Briefly, endothelial cells were grown to confluence on gelatin-coated inserts (0.4-μm polycarbonate membranes) of Transwell Permeable Supports (CoStar Group, Washington DC, USA). After culturing for 5 days, cells on the inserts were incubated with 10 μM dopamine for 30 min, treated with 10 ng/ml VEGF for 90 min, and probed with 1 mg/ml 40-kDa FITC-dextran (MilliporeSigma) for the last 60 min. The amount of FITC-dextran in the lower chamber was measured with a microplate spectrofluorometer (Molecular Devices, Sunnyvale, CA, USA).

### Measurement of endothelial cell apoptosis

Apoptosis of endothelial cells was assessed by TUNEL assay as previously described (Lee et al., 2019). Briefly, cells were fixed with 1% (w/v) paraformaldehyde and 70% (v/v) ethanol on ice and stained with an APO-BrdU TUNEL assay kit (BD Biosciences, San Jose, CA, USA) according to the manufacturer’s protocol and with 1 μg/mL DAPI for 10 min. Apoptotic cells were visualized by confocal microscopy and expressed as the percentage of TUNEL-positive cells.

### Western blot analysis

Lysates from HRECs and mouse retinas were separated by 12% SDS-PAGE and transferred to polyvinylidene fluoride membranes (Lee et al., 2019). The membranes were then incubated with antibodies against tyrosine hydroxylase (1:1000, MilliporeSigma), phospho-Drp1 (Ser616) (1:1000, Cell Signaling Technology, Danvers, MA, USA), mitochondrial fission factor (MFF) (1:1000, Cell Signaling Technology,), Bcl-2-associated X protein (BAX) (1:1000, Cell Signaling Technology), cytochrome c (1:500, Cell Signaling Technology), or β-actin (1:2000, Cell Signaling Technology), followed by incubation with a horseradish peroxidase-conjugated secondary antibody. Protein bands were visualized using a ChemiDoc (Bio-Rad, Hercules, CA, USA).

### Generation of diabetic mice and treatment of mice with insulin and L-dopa

Six-week-old male C57BL/6 mice were obtained from DBL (EumSeong, Korea). The mice were maintained under pathogen-free conditions in a temperature-controlled room with a 12-h light/dark cycle. Diabetic mice were generated by single daily intraperitoneal injections over 5 consecutive days of streptozotocin (50 mg/kg body weight; MilliporeSigma) freshly prepared in 100 mM citrate buffer (pH 4.5), as previously described (Seo et al., 2020). Mice with fasting blood glucose concentrations ≥19 mM and polyuria were considered diabetic. For insulin supplementation, 12 weeks after the first streptozotocin injection, diabetic mice were anesthetized with 3% isoflurane and implanted with an Alzet Mini-Osmotic Pump 2004 (Durect, Cupertino, CA, USA), which delivered human recombinant insulin (MilliporeSigma) at a rate of 58.4 pmol/min/kg. At 18 weeks, a new insulin pump was implanted to maintain normal blood glucose levels for 12 weeks. For L-dopa supplementation, mice were anaesthetized with 3% isoflurane and injected intravitreally with 2 μL of 10 mmol/L L-dopa every two days for 12 weeks. All animal experiments conformed to the *Guide for the Care and Use of Laboratory Animals* (National Institutes of Health, Bethesda, MD, USA) and were approved by the Institutional Animal Care and Use Ethics Committee of Kangwon National University.

### Measurement of VEGF levels in mouse retinas

VEGF levels in mouse retinas were determined using the Quantikine mouse VEGF ELISA kit (R&D Systems, Minneapolis, MN, USA) as previously described (Jeon et al., 2019). Lysates were extracted from the retinas of normal, diabetic, and insulin-supplemented diabetic (HGM) mice, and VEGF levels were determined using a microplate spectrophotometer (Molecular Devices, Sunnyvale, CA, USA) according to the manufacturer’s protocol.

### Measurement of *in vivo* TGase activity and vascular leakage in mouse retinas

*In vivo* TGase activity in the mouse retinas was determined by confocal microscopy as previously described (Lee et al., 2018). Briefly, mice were anaesthetized with 3% isoflurane, and 48 μl of 100 mM 5-(biotinamido)pentylamine was injected into the left ventricle. Enucleated eyes were fixed with 4% paraformaldehyde for 45 min. The retinas were then whole-mounted on glass slides in the Maltese cross configuration and incubated with FITC-conjugated streptavidin (1:200, v/v) for 1 h. *In vivo* TGase activity was visualized by confocal microscopy and quantified by measuring the fluorescence intensities.

Microvascular leakage in the retinas was investigated using fluorescein angiography (Lee et al., 2018). Briefly, mice were anesthetized, and 1.25 mg 500 kDa FITC-dextran (MilliporeSigma) was injected into the left ventricle. Enucleated eyes were fixed with 4% paraformaldehyde, and whole-mounted retinas were visualized by confocal microscopy (Nanoscope Systems). Vascular leakage was quantitatively analyzed by measuring the fluorescence intensities.

### Miles vascular permeability assay in mouse ears and retinas

Miles vascular permeability assay in mouse ears was performed as previously described (Lim et al., 2014). Briefly, Evans blue solution (45 mg/kg, MilliporeSigma) was intravenously injected into the tail vein and allowed to circulate for 30 min. The mice were then anaesthetized and injected intradermally in the middle part of the ears with 10 μL of either 3 ng/μl VEGF or a combination of 3 ng/μl VEGF and 1.9 ng/μl dopamine in PBS. PBS alone was injected into ears as a control. After 30 min, the mice were killed by cervical dislocation and dissected ears were incubated overnight in formamide at 65℃. For the Miles assay in mouse eyes, 4 h after Evans blue injection, retinas were isolated and incubated overnight in formamide at 65℃. The amount of extravasated Evans blue was measured by spectrophotometry (Molecular Device).

### Measurement of ROS generation and ΔΨ_m_ in mouse retinas

ROS levels in mouse retinas were determined using dihydroethidium (Thermo Fisher Scientific) (Lee et al., 2018). Mouse eyes were enucleated and quickly frozen in OCT compound (Sakura Finetek USA, Torrance, CA, USA). Unfixed retinal tissues were cut with a microtome-cryostat (Leica Biosystems, Wetzlar, Germany) at 10 μm thickness, and the sections were stained at 37℃ with 5 μmol/L dihydroethidium in serum-free M199 medium for 30 min. The sections were also incubated with 5 μmol/L MitoSOX^TM^ red (Thermo Fisher Scientific) to measure mitochondrial ROS levels in the retina. The ROS levels were visualized by confocal microscopy (Nanoscope Systems) and quantified by measuring the fluorescence intensities.

To measure ΔΨ_m_ in the retinas, the retinal sections were stained at 37℃ with 2 μmol/L JC-1 in serum-free M199 medium for 30 min. The ΔΨ_m_ was visualized by confocal microscopy and quantified by measuring the fluorescence intensities.

### Visualization of tyrosine hydroxylase expression in mouse retinas

Expression of tyrosine hydroxylase was visualized in sections of mouse retinas by immunohistochemistry. Mouse eyes were fixed with 4% paraformaldehyde, transferred to 30% sucrose, incubated at 4℃ overnight, and frozen in OCT compound. The eyes were then cut with a microtome-cryostat (Leica Biosystems) at 10 μm thickness. The sections were permeabilized with 1% Triton X-100 and stained overnight with anti-tyrosine hydroxylase polyclonal antibody (1:200, MilliporeSigma) at 4℃. The sections were probed with an FITC-conjugated goat anti-rabbit antibody (1:300, MilliporeSigma) for 2 h and stained with 1 μg/mL DAPI for 10 min at room temperature.

### Visualization of VE-cadherin and pericytes in mouse retinas

VE-cadherin in mouse retinas was visualized by confocal microscopy as previously described (Lee et al., 2016). Following whole-mounting in the Maltese cross configuration, the retinas were delipidated with ice-cold acetone for 3 min at -20℃ and permeabilized with 1% Triton X-100 for 4 h. The retinas were then incubated overnight at 4℃ with a monoclonal VE-cadherin antibody (1:100, BD Biosciences) and probed for 2 h with an FITC-conjugated goat anti-rat antibody (1:300, Thermo Fisher Scientific). VE-cadherin in the superficial vascular plexus and deep capillary plexus was visualized in the retinas by confocal microscopy as illustrated in **Fig. 6A**.

To visualize pericytes, whole-mounted retinas were incubated overnight at 4℃ with a monoclonal neuron-glial antigen 2 (NG2) antibody (1:200, MilliporeSigma) and probed for 2 h with an FITC-conjugated goat anti-rabbit antibody (1:300) and the endothelial cell-specific marker Alexa 647-Isolectin B4 (1:1000; Thermo Fisher Scientific). The number of NG2-positive pericytes was counted in the superficial vascular plexus of the retina per mm^2^ vascular area.

### Retinal trypsin digestion assay and apoptosis of retinal vasculature

Retinal vasculature was isolated by trypsin digestion as previously described (Chou et al., 2013). Briefly, mouse retinas were isolated from eyes fixed in 10% neutral buffered formalin (MilliporeSigma) and incubated for 3 h with 3% trypsin in 0.1 M Tris buffer (pH 7.8) at 37℃. The digested retinas were flat-mounted on glass slides, stained with Periodic Acid Schiff (MilliporeSigma), and observed under a light microscope. The numbers of acellular capillaries and pericyte ghosts were counted per mm^2^ vascular area.

To assess apoptosis in the retinal vasculature, flat-mounted digested retinas were stained with the APO-BrdU TUNEL assay kit (BD Biosciences) and incubated for 1 h with Alexa Fluor 647-Isolectin B4 (Thermo Fisher Scientific). TUNEL-positive cells in the stained retinas were visualized using confocal microscopy.

### Statistical analysis

Data were analyzed using OriginPro 2015 software (OriginLab, Northampton, MA, USA). Data are expressed as means ± standard deviation (SD) of at least three independent *in vitro* experiments or six independent *in vivo* experiments. Statistical significance was determined using one-way ANOVA with Holm-Sidak’s multiple comparisons test. *P* values of < 0.05 were considered statistically significant.

## Author Contributions

Y.-J.L. designed and performed experiments, analyzed data, and wrote the manuscript. H.- Y.J., A.-J.L., and M.K. designed experiments and analyzed data. K.-S.H. conceptualized the study, designed experiments, analyzed and interpreted data, and wrote the manuscript. All authors approved the final version of the manuscript.

